# Ancient hybridization leads to the repeated evolution of red flowers across a monkeyflower radiation

**DOI:** 10.1101/2023.02.13.528387

**Authors:** Aidan W. Short, Matthew A. Streisfeld

## Abstract

The re-use of old genetic variation can promote rapid diversification in evolutionary radiations, but in most cases, the historical events underlying this divergence are not known. For example, ancient hybridization can generate new combinations of alleles that sort into descendant lineages, potentially providing the raw material to initiate divergence. In the *Mimulus aurantiacus* species complex, there is evidence for widespread gene flow among members of this radiation. In addition, allelic variation in the *MaMyb2* gene is responsible for differences in flower color between the closely related ecotypes of subspecies *puniceus*, contributing to reproductive isolation by pollinators. Previous work suggested that *MaMyb2* was introgressed into the red-flowered ecotype of *puniceus*. However, additional taxa within the radiation have independently evolved red flowers from their yellow-flowered ancestors, raising the possibility that this introgression had a more ancient origin. In this study, we used repeated tests of admixture from whole-genome sequence data across this diverse radiation to demonstrate that there has been both ancient and recurrent hybridization in this group. However, most of the signal of this ancient introgression has been removed due to selection, suggesting that widespread barriers to gene flow are in place between taxa. Yet, a roughly 30 kb region that contains the *MaMyb2* gene is currently shared among the red-flowered taxa. Patterns of admixture, sequence divergence, and extended haplotype homozygosity across this region confirm a history of ancient hybridization, where functional variants have been preserved due to positive selection in red-flowered taxa but lost in their yellow-flowered counterparts. The results of this study reveal that selection against gene flow can reduce genomic signatures of ancient hybridization, but that historical introgression can provide essential genetic variation that facilitates the repeated origins of phenotypic traits between lineages.

## Introduction

The rapid diversification rates and extraordinary levels of phenotypic variation found in evolutionary radiations provide excellent opportunities to study the processes of adaptation and speciation. Many classic models suggest that adaptation and reproductive isolation evolve due to the accumulation of new mutations within isolated lineages (Orr 1995;1998, Barton 2001). However, the extensive phenotypic diversity found in radiations often arises too quickly to be explained by the long waiting times between new mutations (Hedrick 2013). Therefore, rapid diversification is likely to depend on pre-existing genetic variation. This variation either can originate in an ancestral population (e.g., Choi et al. 2021), or it can be transferred across species boundaries during divergence due to natural hybridization and introgression (e.g., Han et al. 2017, Meier et al. 2017). Indeed, the recognition that gene flow is common during evolutionary radiations has led some to propose that introgressive hybridization may be an important contributor of rapid speciation (Marques et al. 2019).

Determining the history of hybridization in evolutionary radiations can help us to understand the role that gene flow has played in generating biodiversity. Recent or ongoing hybridization between lineages can lead to the transfer of adaptive genetic variation that directly contributes to reproductive isolation between diverging populations (Pardo-Diaz et al. 2012). Alternatively, hybridization can occur much earlier in a radiation, which can facilitate rapid and repeated diversification (Meier et al. 2019). The re-use of old genetic variation that pre-dates species splitting events immediately generates large amounts of polymorphism in populations, which can lead to novel phenotypes and drive adaptive divergence and reproductive isolation (Marques et al 2019). In addition, introgressed alleles will tend to be at higher frequencies than new mutations, making them less likely to be lost to drift (Barrett and Schluter 2008). Moreover, when these events coincide with novel ecological opportunity, rapid diversification becomes possible. For example, hybridization early in the divergence history of African cichlid fishes (Meier et al. 2017), Darwin’s finches (Han et al. 2017), and the Hawaiian silverswords (Barrier et al. 1999) has played a critical role in the rapid diversification of these radiations.

Despite the potential for hybridization to lead to rapid diversification, in most cases, introgressed genetic variation will be neutral or deleterious when shared between species (Mallet 2005). Deleterious effects arise because of genetic incompatibilities that occur in the recipient’s genetic background or due to maladaptation of the transferred genetic material in a divergent ecological environment (Dobzhansky 1982, Nosil 2012). Natural selection operating against deleterious introgressed ancestry will result in its removal from populations, the extent of which will be modulated by variation in the local recombination rate (Schumer et al. 2018). Specifically, selection will be highly efficient at removing blocks of introgressed ancestry if they occur in regions of low recombination, as deleterious alleles are less likely to be separated from neutral or beneficial variation (Brandvain et al. 2014, Aeschbacher et al. 2017). This is expected to be more pronounced under highly polygenic architectures of reproductive isolation, as more of the genome acts as a barrier to gene flow (Martin et al 2019). As a result, selection against gene flow will result in a heterogeneous pattern of introgression across the genome (Ellegren et al. 2012, Nelson et al. 2021, Liu et al. 2022).

Repeated phenotypic transitions within recent radiations provide excellent opportunities to study the evolutionary history of the genetic variation responsible for adaptive trait differences. Over the past decade, an increasing amount of evidence suggests that introgressive hybridization can be an important source of the genetic variation used for repeated trait evolution (e.g. De-Kayne et al. 2022). In this study, we take advantage of the independent evolution of red flower color among the members of the *Mimulus aurantiacus* species complex to determine the genome-wide effects and evolutionary history of hybridization that may have contributed to adaptation and speciation within this radiation.

The *Mimulus aurantiacus* species complex (Phrymaceae) consists of seven closely related, woody shrub subspecies that radiated across California after they diverged from their sister species *M. clevelandii* roughly one million years ago (Stankowski et al 2019). The subspecies inhabit a diverse range of environments and show remarkable phenotypic differentiation, primarily in their flowers (Fig. 1) (Chase et al. 2017). However, all of the taxa are interfertile to varying degrees (McMinn 1951), and there is evidence for gene flow between many of them (Stankowski et al. 2019). However, the specific patterns and history of gene flow among the members of this complex and their role in reproductive isolation have not been explored.

**Figure 1.**
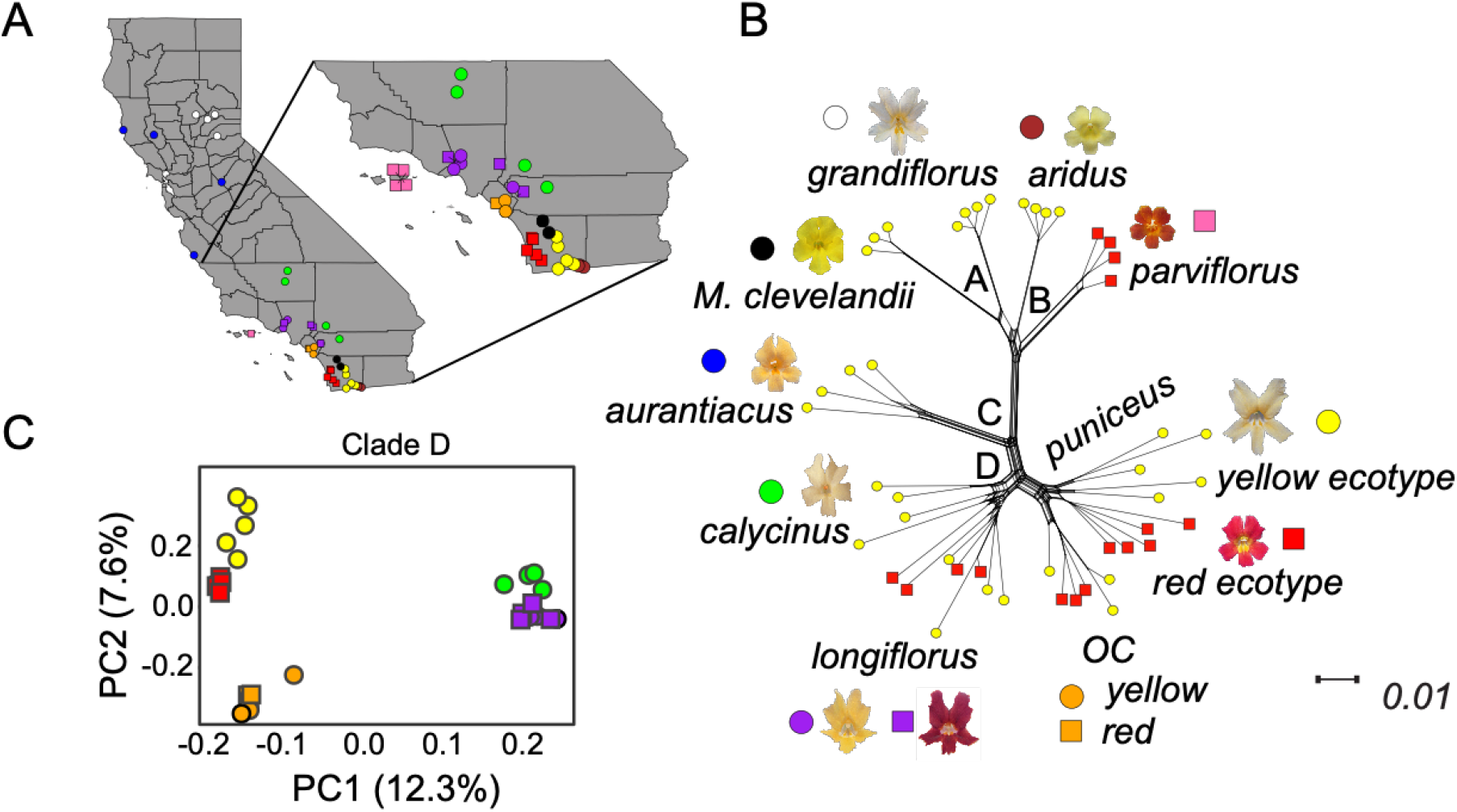
Geographic sampling and patterns of relatedness among members of the *M. aurantiacus* species complex. (A) Locations of the samples used for whole genome sequencing from the seven subspecies of the *Mimulus aurantiacus* species complex and the outgroup, *M. clevelandii* across California. Colors correspond to the different taxa, with *M. clevelandii* in black, *grandiflorus* in white, *aridus* in brown, *aurantiacus* in blue, *calycinus* in green, *longiflorus* in purple, *parviflorus* in pink, Orange County *puniceus* in orange, the red ecotype of *puniceus* from San Diego in red and the yellow ecotype of *puniceus* from San Diego in yellow. Circles correspond to plants with yellow flowers and squares correspond to plants with red flowers. (B) Splitstree network showing the evolutionary relationships of the taxa, with representative photographs of their flowers. The four major clades of the radiation are labelled with black letters. Symbols next to each photograph match those in panel A. Tips of the tree are colored according to the flower color of the individual plant, with yellow circles denoting yellow flowers and red squares corresponding to red flowers. (C) Plot of the first two principal component axes from taxa in Clade D. The percent variation explained by each axis is reported. Shapes and colors match those in panel A.

Most work in this complex has focused on the red- and yellow-flowered ecotypes of subspecies *puniceus*, which display a phenotypic transition along a west to east gradient in San Diego County (hereafter, referred to as the red and yellow ecotypes, respectively). The primary trait that distinguishes the ecotypes is flower color, with allelic variation in the *MaMyb2* gene largely responsible for the derived, anthocyanin pigment found in the flowers of the red, but not yellow, ecotype (Streisfeld et al 2013). Divergence between these ecotypes is maintained by pollinator-mediated selection, with hummingbirds preferring red flowers and hawkmoths preferring yellow flowers (Streisfeld and Kohn 2007, Sobel and Streisfeld 2015). Moreover, introgression of *MaMyb2* from a distantly related subspecies in the complex led to the formation of the novel, red-flowered phenotype that was influential for pollinator-mediated reproductive isolation (Stankowski and Streisfeld 2015).

However, despite the importance of this novel phenotype in the red ecotype, transitions from yellow to red flowers also occurred in other lineages in this radiation. For example, there is a rare, red-colored form that occurs in some populations of the predominantly yellow-flowered subspecies *longiflorus*. In addition, red and yellow flowers are found in a separate population series of subspecies *puniceus* to the north of San Diego (Beeks 1962) (Figure 1). The red variant of *longiflorus* was described previously as *M. rutilus*, but genetic data reveal that it is nearly indistinguishable from the yellow-flowered form of *longiflorus* (Chase et al. 2017). There is little known about the red- and yellow-flowered variants of *puniceus* to the north, but Beeks (1962) used morphological evidence to suggest that northern and southern *puniceus* should be divided into two population series. Similarly, Streisfeld and Kohn (2005) found that northern and southern *puniceus* were genetically differentiated at two chloroplast DNA markers. The presence of red flowers in these distinct populations of *puniceus* and *longiflorus* thus raises questions about the origin of the genetic variation responsible for red flowers. Specifically, is the same variant shared among these taxa, which could suggest that hybridization deeper in their evolutionary history may have fueled the repeated origin of this diversity? Or have there been independent genetic origins of red flowers in these groups?

In this study, we examined the history and genomic consequences of introgression within this radiation to reveal its impact on the repeated origins of red flowers. We identified evidence of both recurrent and ancient hybridization between lineages, but extensive selection against gene flow has erased much of the genomic signal of this introgression. Nevertheless, we found that introgression was responsible for the transfer and subsequent maintenance of a common haplotype in the *MaMyb2* gene between lineages, where it led to the repeated evolution of red flowers. These results reveal that the accumulation of reproductive barriers between divergent taxa can reduce genomic signatures of ancient hybridization, but introgression can still provide the functional variation that facilitates the repeated origins of phenotypic traits and adaptive divergence between lineages.

## Results and discussion

### Multiple transitions from yellow to red flowers among lineages

Previous phylogenetic analyses using both reduced representation and whole genome sequencing revealed four monophyletic clades that delineated the members of the *M. aurantiacus* species complex (Stankowski and Streisfeld 2015, Chase et al 2017, Stankowski et al 2019). Clade A consisted entirely of individuals of *M. a. ssp. grandiflorus* from northeastern California. Clade B included individuals from *M. a. ssp. aridus* from southeastern California and *M. a. ssp. parviflorus* that is endemic to the Channel Islands off the California coast. Clade C comprised samples from *M. a. ssp. aurantiacus* in central and northern California. The highly diverse Clade D from southern California included *M. a. ssp. calycinus, M. a. ssp. longiflorus*, and the red and yellow ecotypes of *M. a. ssp. puniceus*. Using 10 more whole genome sequences generated here (Table S1), we inferred patterns of relatedness among additional red-flowered samples. Specifically, we sequenced four individuals from the red-flowered *M. a. ssp. longiflorus* and three individuals each from red and yellow-flowered populations of the northern *M. a. ssp. puniceus* collected from Orange County. These newly generated sequences were combined with 37 whole genomes from the 7 subspecies and 2 ecotypes (*n* = 4–5 per taxon) and their sister taxon *M. clevelandii* (*n* = 3) that were described in Stankowski et al. (2019). After reads were aligned to the *M. aurantiacus* reference genome, we performed variant calling, filtering, and haplotype phasing as described previously (Stankowski et al 2019), which resulted in a final dataset containing 12,730,568 SNPs. Hereafter, we refer to the taxa only by their subspecies name.

To determine the relationships among the additional red-flowered forms, we generated a neighbor-joining splits network that included all 47 samples. The network revealed patterns consistent with previous analyses that defined the four primary clades (Fig. 1B). The relationships among the 27 samples from clade D were also confirmed using principal components analysis (PCA), which revealed five groups that corresponded to the five taxa of clade D referred to in this study (Fig. 1C). Of note, these analyses revealed a clear separation between the northern and southern *puniceus* populations, suggesting that *puniceus* from Orange County is a distinct lineage that likely diverged prior to the split between the red and yellow ecotypes. In addition, even though the Orange County *puniceus* contains populations with either red or yellow flowers, the splits network and the PCA did not identify separation between flower color morphs from Orange County. Thus, the red and yellow forms of the northern *puniceus* do not appear to be as diverged as the red and yellow ecotypes in San Diego, but additional work is needed to quantify levels of reproductive isolation between these forms. In addition, the red and yellow forms of *longiflorus* were interdigitated with each other (Fig 1B), confirming that the red-flowered variants represent a polymorphism within *longiflorus*, rather than a distinct taxonomic entity. Hereafter, the samples from the northern *puniceus* will be referred to as *OC*, for Orange County *puniceus*.

These findings reveal that the red variants in *longiflorus* and *OC puniceus* represent additional, independent transitions to red flowers from each of their yellow-flowered ancestors. Specifically, the red variants of *longiflorus* and *puniceus* do not group with the other red-flowered taxa (*parviflorus* or the red ecotype), consistent with previous ancestral state reconstructions that indicated the independent derivation of red flowers (Stankowski and Streisfeld 2015). However, all red-flowered taxa produce anthocyanin pigments in their flowers, and the red ecotype and *parviflorus* share a common genetic basis controlling the production of red flowers (Stankowski and Streisfeld 2015). Thus, these results suggest either repeated origins of red flowers, or that there was a single origin that occurred deep in the evolutionary history of clade D, followed by multiple losses.

### Genome-wide evidence of gene flow

Although there is evidence that red flowers arose in the red ecotype due to prior introgression of *MaMyb2* with *aridus* (Stankowski and Streisfeld 2015), there is currently no information on the genomic extent of this gene flow between these taxa. Moreover, these two taxa do not currently come into geographic contact with one another. By contrast, the yellow ecotype does contact populations of *aridus* in certain parts of its range (Thompson 2005), suggesting that *aridus* may have hybridized with the yellow ecotype more recently than the red ecotype. We tested for genome-wide evidence of introgression among these taxa using *D*-statistics (Green et al. 2010). Patterson’s *D* measures asymmetry between the numbers of sites with ABBA and BABA patterns (where A and B are ancestral and derived alleles, respectively) across a phylogeny with three ingroup taxa and an outgroup that have the relationship (((P1, P2), P3) O). A significant excess of either pattern gives a nonzero value of *D*, which is taken as evidence that gene flow has occurred between P3 and one of the ingroup taxa.

We calculated Patterson’s *D*-statistic among all possible triplets of ingroup taxa (Table S2), but we focus here on differences that occurred when the red or yellow ecotypes were set as P2. Specifically, we set different members of clades C and D as P1, the red or yellow ecotypes as P2, *aridus* as P3, and *M. clevelandii* as the outgroup. We found that the yellow ecotype displayed greater genome-wide evidence of introgression with *aridus* relative to the red ecotype. This is indicated by a higher value of Patterson’s *D*-statistic when the yellow ecotype was P2, regardless of which taxon was used as P1 (Table 1). This suggests that more extensive or more recent introgression has occurred between *aridus* and the yellow ecotype compared to the red ecotype. By contrast, even though *parviflorus* also has red flowers, there was no significant evidence for introgression when we repeated these tests using *parviflorus* as P3 (Table S2). These findings are consistent with the current geographic distribution of *parviflorus*, which is endemic to the Channel Islands off the coast of California and is isolated from the taxa on the mainland.

**Table 1.** *D*-statistics, with associated Z-scores and p-values calculated using either the red or yellow ecotypes as P2, *aridus* as P3, *M. clevelandii* as the outgroup, and various taxa from clades C and D as P1.

| P1 | P2 | P3 | *D*-Statistic | Z-score | p-value |
| --- | --- | --- | --- | --- | --- |
| *aurantiacus* | red ecotype | *aridus* | 0.072 | 5.571 | $2.5 \times 10^{-8}$ |
| *calcyinus* | red ecotype | *aridus* | 0.052 | 4.889 | $1.0 \times 10^{-8}$ |
| *longiflorus* | red ecotype | *aridus* | 0.074 | 6.305 | $2.9 \times 10^{-10}$ |
| *OC* | red ecotype | *aridus* | 0.044 | 6.578 | $4.8 \times 10^{-11}$ |
| *aurantiacus* | yellow ecotype | *aridus* | 0.098 | 7.929 | $2.2 \times 10^{-15}$ |
| *calcyinus* | yellow ecotype | *aridus* | 0.082 | 7.924 | $2.3 \times 10^{-15}$ |
| *longiflorus* | yellow ecotype | *aridus* | 0.103 | 9.502 | 0 |
| *OC* | yellow ecotype | *aridus* | 0.074 | 11.646 | 0 |
| red ecotype | yellow ecotype | *aridus* | 0.035 | 5.366 | $8.1 \times 10^{-8}$ |

### Ancient and recurrent hybridization throughout the history of this radiation

Although powerful for detecting genome-wide evidence of gene flow, the *D*-statistic can only identify introgression that has occurred after the two sister taxa used in the test have diverged from each other. This is because genetic variation that was transferred into the ancestor of these sister taxa would be inherited by both modern-day taxa, resulting in them both sharing alleles with the third taxon (Fig. 2A). By taking advantage of the diversity of taxa in the *M. aurantiacus* species complex, we ran multiple tests for introgression using successively more distantly related taxa as P1, which allowed us to identify allelic differences that were shared between the yellow ecotype (P2) and *aridus* (P3) that occurred after the yellow ecotype split with the P1 taxon (Fig. 2). Thus, by gradually increasing the phylogenetic distance between the two sister taxa used in these tests, we should be able to identify introgression that occurred at various points further back in time.

**Figure 2:**
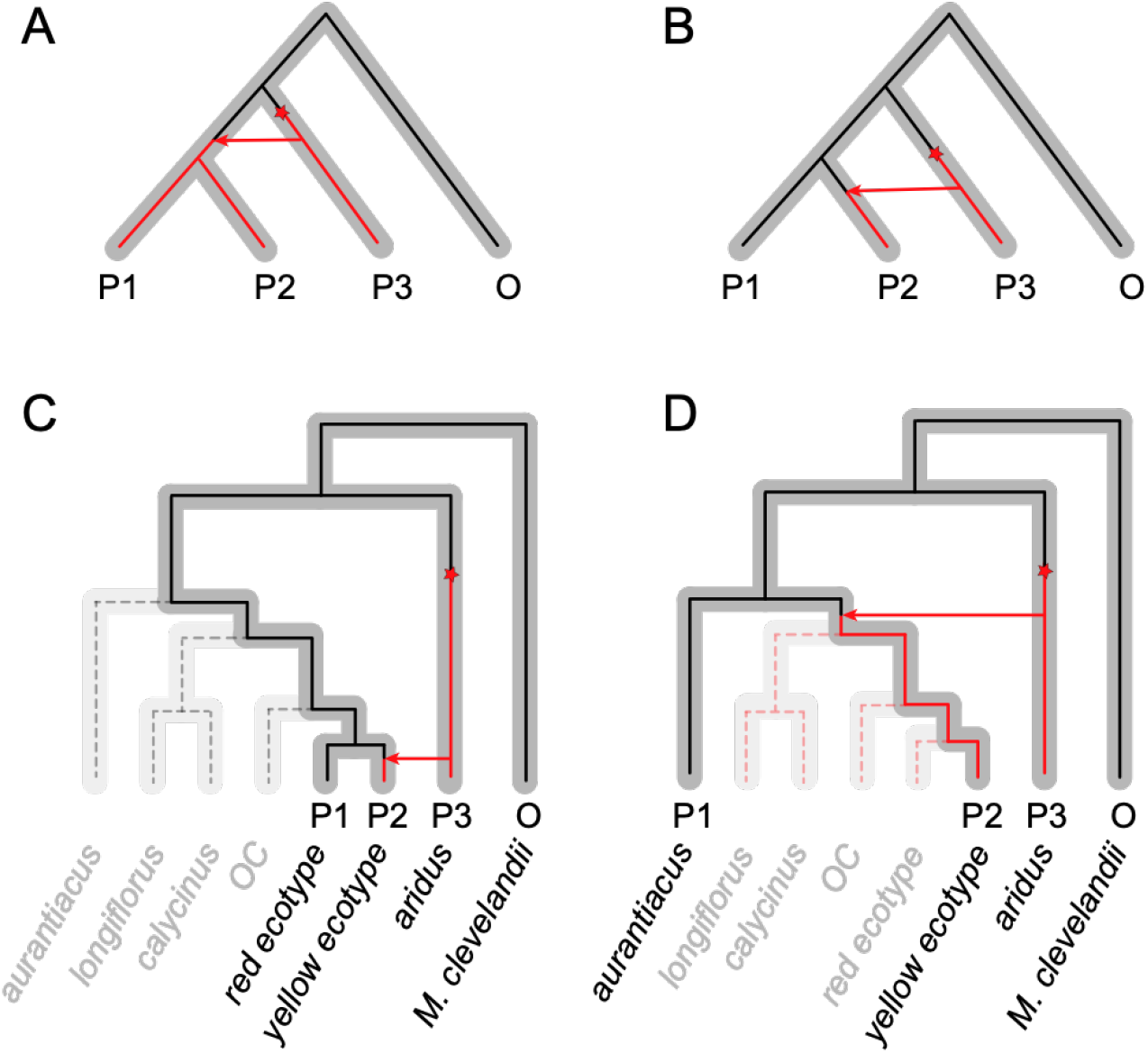
Evolutionary radiations can help evaluate the history of introgression. (A-B) In a four taxon tree, with three ingroups (P1-P3) and an outgroup (O), introgression between P2 and P3 can only be identified if gene exchange occurred after the split between the two sister taxa (P1 and P2). The species trees are in gray, and the ancestral (black) and derived (red) alleles at a locus are indicated. (A) A mutation (red star) that occurs in the lineage leading to P3 that is transferred via introgression into an ancestor of P1 and P2 will result in the sharing of that genetic information in both P1 and P2. As a consequence, this deeper history of introgression will be undetectable using tests, such as *D*-statistics. (B) However, introgression between P2 and P3 can be detected if the genetic exchange occurred after P1 and P2 diverged from each other, as P1 would retain the ancestral sequence, but P2 would contain the derived variant that is shared with P3. (C-D) The species tree from the *M. aurantiacus* radiation is presented in gray, but tips are grayed out to show only the four taxa used in the tests of introgression (denoted by P1-P3, and O). By using different P1 taxa at increasing levels of divergence with the yellow ecotype, we can track introgression that occurred deeper in time. (C) Only genetic variation that was introgressed after the divergence of the red and yellow ecotypes of *puniceus* can be identified when the red ecotype is used as P1 and the yellow ecotype is used as P2. The black dashed line represents the retention of the ancestral allele in the taxa not being used in the test. (D) When a more distant taxon, such as *aurantiacus*, is used as P1, genetic variation that was introgressed in all of clade D can be identified.

To obtain estimates of the variation in introgression along the genome, we calculated (*f*_*d*_) in 50 kb windows. Patterson’s *D*-statistic is intended for genome-wide estimates of gene flow, but *f*_*d*_ provides a measure of the admixture proportion that has been modified for use in genomic windows (Martin et al 2015). We calculated *f*_*d*_ repeatedly, each time using one of the five taxa that occurred at different levels of sequence divergence from the yellow ecotype as P1. In all tests, the yellow ecotype was used as P2, *aridus* was used as P3, and *M. clevelandii* was the outgroup. This allowed us to determine how levels of introgression varied across the genome at different periods throughout the history of this radiation. Consistent with our calculations of Patterson’s *D*-statistic (Table 1), the mean *f*_*d*_ among windows was greater than zero in all tests, regardless of which taxon was used as P1. Qualitatively similar results were found when the red ecotype was set as P2. This suggests that introgression with *aridus* or its ancestor has occurred at multiple points throughout the divergence history of clade D. In addition, gene flow increases significantly with levels of sequence divergence between the P1 taxon and the yellow ecotype (measured as *d*_*a*_), with mean *f*_*d*_ nearly doubling when *aurantiacus* is P1 compared to when the red ecotype is P1 (Fig. 3, Table S3). This provides clear evidence that introgression dates back to at least the ancestor of clade D after it diverged from clade C. Although additional taxonomic diversity would be needed to determine if gene flow occurred even further back in the radiation, these results reveal the historical presence of hybridization between lineages.

**Figure 3.**
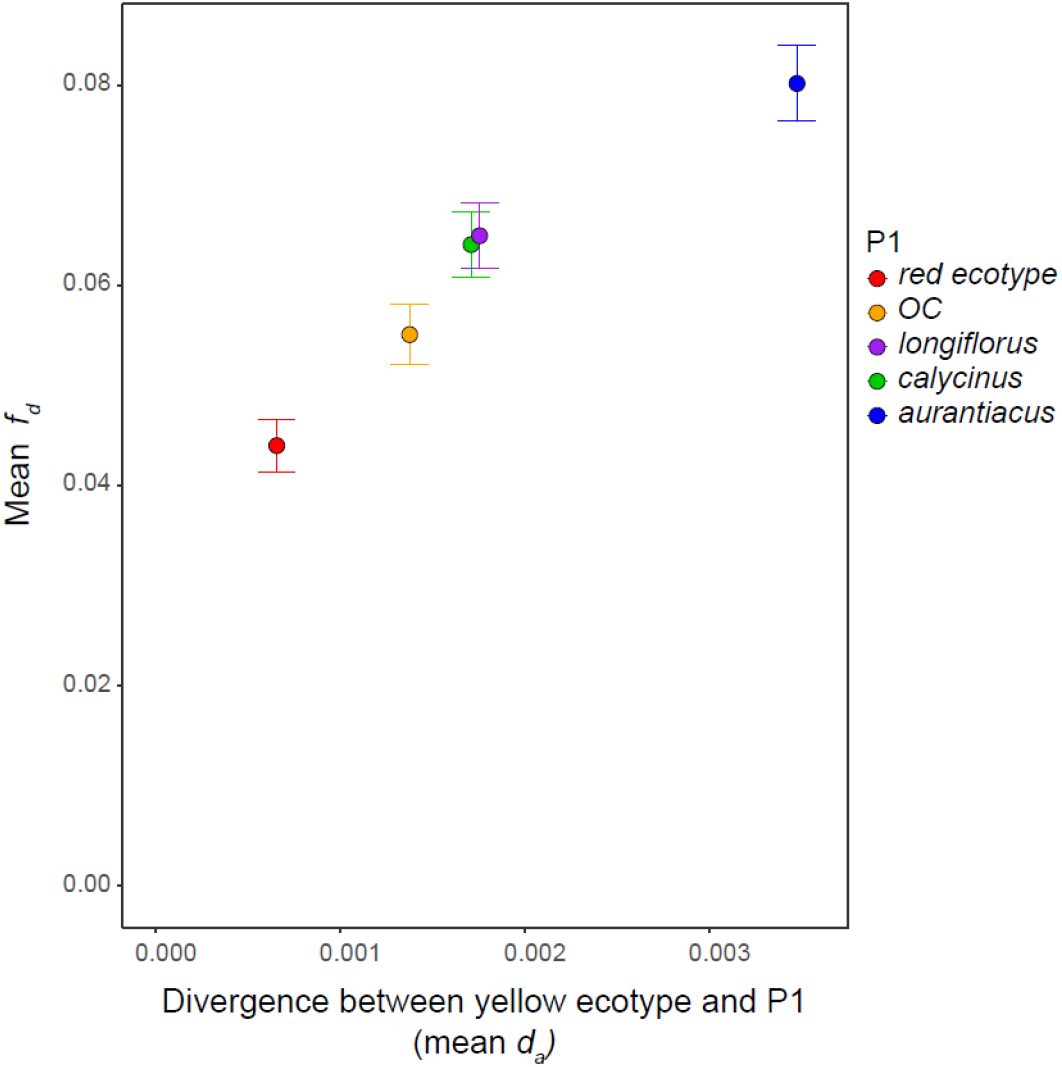
A long history of gene flow in the radiation. Mean and 95% confidence intervals of the admixture proportion (*f*_*d*_) calculated in 50 kb windows are plotted against mean levels of sequence divergence (*d*_*a*_) between the yellow ecotype and taxa in clades C and D. *f*_*d*_ was calculated repeatedly in each window, with one of five different taxa as P1, the yellow ecotype as P2, *aridus* as P3, and *M. clevelandii* as the outgroup. The increase in mean *f*_*d*_ with divergence time indicates that admixture with the lineage leading to *aridus* has occurred throughout the radiation, and as far back as the ancestor of clade D.

### Widespread selection against gene flow has removed much of the signal of introgression

To determine the genomic consequences of this hybridization, we plotted how the admixture proportion (*f*_*d*_) varied across the genome. This revealed extensive heterogeneity in introgression (Fig. 4A), with broad regions of elevated *f*_*d*_ on each chromosome that were interspersed with regions of little to no admixture. This same pattern was found regardless of which taxon was set as P1, though the peaks of *f*_*d*_ tended to vary in height based on levels of divergence with the yellow ecotype (see also Fig. 3). Moreover, we found that the distribution of *f*_*d*_ values among windows was highly skewed, with much of the density near zero (Fig. S1), indicating that despite a few regions where introgression has been maintained in the genome, little of the signal of introgression remains.

**Figure 4.**
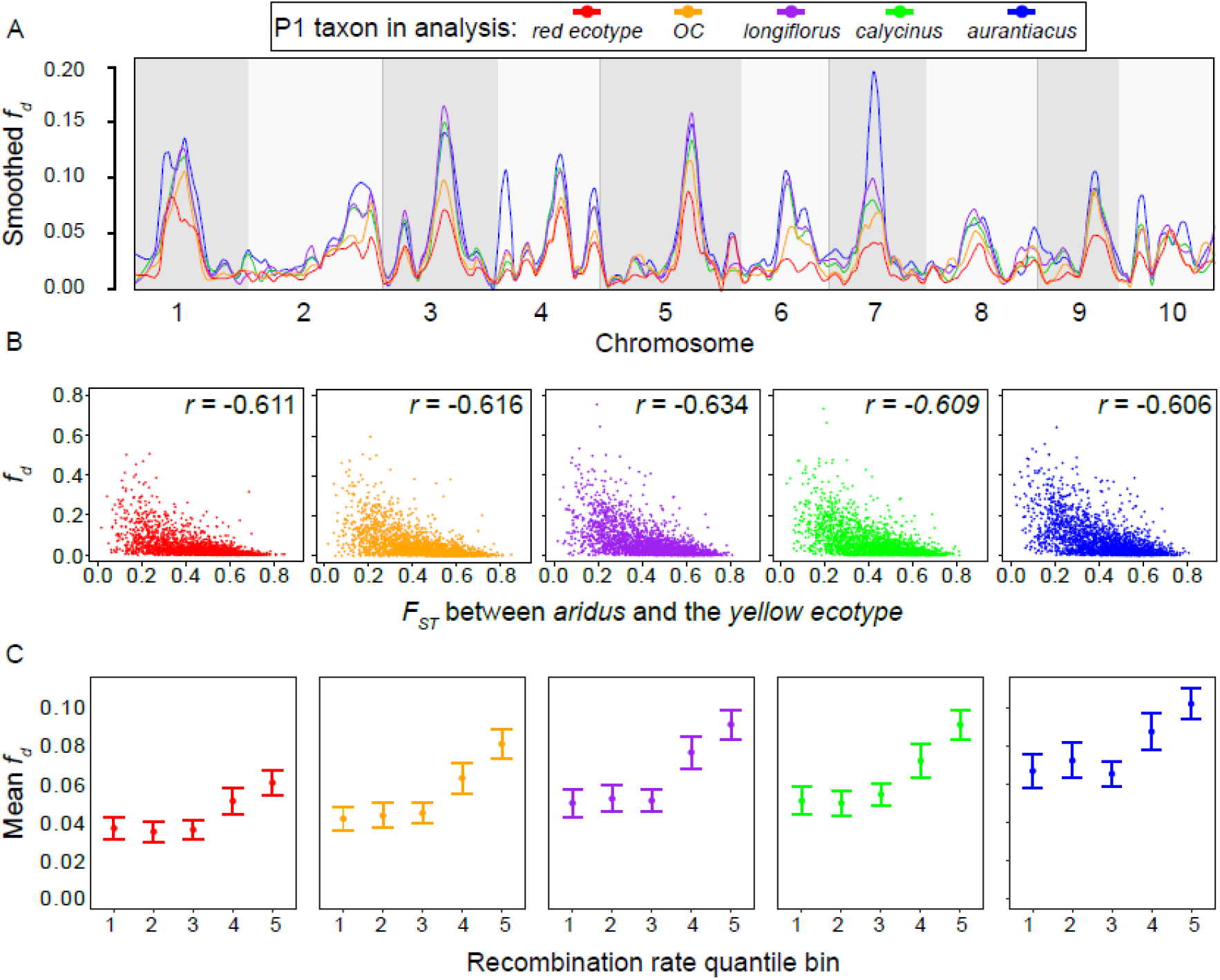
Heterogeneous introgression across the genome indicates selection against gene flow. (A) Loess-smoothed *f*_*d*_ values plotted across the 10 chromosomes of the *Mimulus aurantiacus* genome, with the yellow ecotype as P2, *aridus* as P3, and one of five different taxa as P1. Colors indicate the taxa from clades C and D used as P1. (B) Scatterplot showing the relationship between *F*_*ST*_ and *f*_*d*_ in these same 50 kb windows, with the correlation coefficient between the statistics in the upper right hand corner of each plot. (C) Mean and 95% confidence intervals of *f*_*d*_ in different quantile bins of recombination rate. Quantile bins of recombination rate are as follows: 1 = 0 – 0.824 cM/Mb; 2 = 0.824 – 1.51 cM/Mb; 3 = 1.51 – 2.33 cM/Mb; 4 = 2.33 – 3.66 cM/Mb; 5 = 3.66 – 10.9 cM/Mb.

Given that the entire genome is admixed initially upon hybridization, it is important to determine why some regions of the genome maintain evidence for introgression and others do not. By characterizing the regions of the genome that fail to introgress between taxa, we potentially can identify loci that act as barriers to gene flow. Specifically, natural selection can result in heterogeneous patterns of introgression across the genome, because introgressed genetic variants that are deleterious will tend to be eliminated from a population following admixture (Ellegren et al. 2012, Nelson et al. 2021, Liu et al. 2022). It is well known that local rates of recombination also can modulate whether introgressed tracts are maintained, particularly as the number of loci responsible for reproductive isolation increases (Brandvain et al. 2014, Aeschbacher et al. 2017, Schumer et al. 2018, Martin et al. 2019). This is because deleterious variants are expected to become decoupled from neutral variants more rapidly in regions of higher recombination, allowing a more pronounced signal of introgression to be maintained. By contrast, introgressed deleterious and neutral variants will remain in linkage disequilibrium in regions of low recombination, which will make it easier for selection to purge introgressed variation from the population (Brandvain et al. 2014, Aeschbacher et al. 2017, Schumer et al. 2018). Thus, under scenarios where a polygenic architecture contributes to reproductive isolation, we expect a heterogeneous pattern of admixture across the genome, such that the admixture proportion is correlated with levels of genetic divergence and recombination rate (Martin et al 2019).

To determine the impacts of selection on genome wide patterns of introgression, we examined the relationship between *f*_*d*_ and genetic differentiation (*F*_*ST*_). *F*_*ST*_ is a relative measure of divergence that reflects differences in allele frequencies between populations (Wright 1950, Weir and Cockerham 1984). Because gene flow opposes divergence, *F*_*ST*_ will be reduced in genomic regions of elevated admixture (Martin et al. 2013). Indeed, the elevated *f*_*d*_ that we detected in certain areas of the genome suggests that there is (or was) gene flow between the yellow ecotype and *aridus*. Given that *f*_*d*_ is proportional to the effective migration rate (Martin et al 2015), we expect that selection against gene flow should result in locally elevated *F*_*ST*_ but reduced *f*_*d*_. Consistent with these predictions, we found a strong negative correlation between *f*_*d*_ and *F*_*ST*_, regardless of which taxon was used as P1, with the strength of the correlation ranging from −0.606 to −0.634 (Fig. 4B). We also found a consistently strong positive correlation between *f*_*d*_ and levels of nucleotide diversity (π) (Fig. S2), which is to be expected given the negative relationship between *F*_*ST*_ and π (Stankowski et al 2019). Combined with evidence of ancient and recurrent introgression between the lineage leading to *aridus* and the ancestor of clade D, the observed relationship between *f*_*d*_ and *F*_*ST*_ (and π) suggests that historical divergence between these lineages has been shaped by multiple episodes of selection generated by the accumulation of numerous isolating barriers. These results are consistent with previous simulations that revealed selection against gene flow as a plausible explanation for the heterogeneous genomic landscapes found in these *Mimulus* taxa (Stankowski et al 2019).

To examine the relationship between recombination rate and introgression, we sorted the *f*_*d*_ values into quantile bins based on estimates of recombination rates calculated in 500 kb windows from Stankowski et al. (2019). We then calculated the mean and 95% confidence intervals for *f*_*d*_ within each recombination rate quantile bin. We found that, regardless of which taxon was set as P1 for the calculation of *f*_*d*_, the two bins with the highest recombination rate (2.33-3.66 cM/Mb and 3.66-10.9 cM/Mb) had significantly higher mean values of *f*_*d*_ than the three bins with lower recombination rates (0-0.824 cM/Mb, 0.824-1.51 cM/Mb and 1.51-2.33 cM/Mb) (Fig. 4C, Table S4). Thus, regions of high recombination appear to maintain greater levels of introgressed genetic variation than regions with low recombination, contributing to the heterogeneous pattern that we detected (Fig. 4A).

In sum, we found a genome-wide negative correlation between *f*_*d*_ and *F*_*ST*_, as well as a positive relationship between *f*_*d*_ and recombination rate (and π), all of which are consistent with widespread selection against gene flow and a polygenic architecture of reproductive isolation due to some combination of hybrid incompatibilities or ecologically-based local adaptation. Although more detailed studies of ecological divergence between these taxa are warranted, reductions in F1 hybrid seed viability and pollen fertility exist in crosses between *aridus* and several members of clade D, indicating the presence of postzygotic barriers (J.M. Sobel, personal communication). Similar patterns have been observed in swordtail fish (Schumer et al. 2018) and *Heliconius* butterflies (Martin et al. 2019), further revealing the importance of polygenic genetic architectures underlying both prezygotic and postzygotic forms of reproductive isolation.

### Ancient hybridization leads to repeated transitions of red flowers

Despite the barriers to gene flow that are likely in place now, the current findings indicate that ancient hybridization was possible between ancestral individuals of *aridus* and members of clade D, thus raising the possibility that the *MaMyb2* allele responsible for red flowers was present in the ancestor of this clade. This conclusion would be supported if the same allele was responsible for the repeated origins of red flowers in the different lineages of clade D. Moreover, we would expect to find common signatures of admixture and genetic divergence among the red-flowered taxa across the genomic region containing *MaMyb2* that does not exist with their closely-related yellow-flowered counterparts. Specifically, we would expect to see elevated *f*_*d*_ but reduced genetic divergence (*d*_*xy*_) when the red-flowered samples are compared to samples from *aridus*. By contrast, we predict that there will be elevated divergence in this region of the genome when red-flowered samples are compared to their most closely related yellow-flowered partners.

We scanned the genomes of these taxa and calculated *f*_*d*_ and *d*_*xy*_ in shorter, overlapping 10 kb windows (step size of 100 bp). We calculated *f*_*d*_ separately for each of the three red-flowered groups in clade D (i.e., red ecotype, red-*OC*, and red *longiflorus*), which were each set as P2, and we set *aurantiacus* as P1 and *aridus* as P3. Across a roughly 30 kb region that contains the *MaMyb2* gene, we see a substantial increase in *f*_*d*_ that is considerably larger than the genome-wide average *f*_*d*_ values for these comparisons, indicating that numerous sites across this region are admixed between *aridus* and the red taxa (Fig. 5A). In addition, *d*_*xy*_ between red-flowered samples from clade D and *aridus* is reduced in this same region relative to their most closely related yellow-flowered partners (i.e., yellow ecotype, yellow-*OC*, and yellow *longiflorus*) (Fig. 5B). However, just downstream of this region, the red- and yellow-flowered samples from clade D all show elevated divergence with *aridus*, which is consistent with the phylogenetic patterns of relatedness between these taxa. These results indicate that this region corresponds to a common block of introgression found in each of the red-flowered taxa that is absent from their yellow-flowered counterparts. The position along the chromosome where this block ends on the 5’ end remains unclear, as divergence is unusually low for all samples. Additional long-read sequencing will likely be necessary to determine the precise borders of this introgressed block. Finally, we detected elevated divergence between each of the red-flowered taxa and their most closely related yellow-flowered partner in this same region (Fig. 5C), consistent with this block of introgression being shared in the red-flowered taxa but not in the yellow-flowered plants.

**Figure 5.**
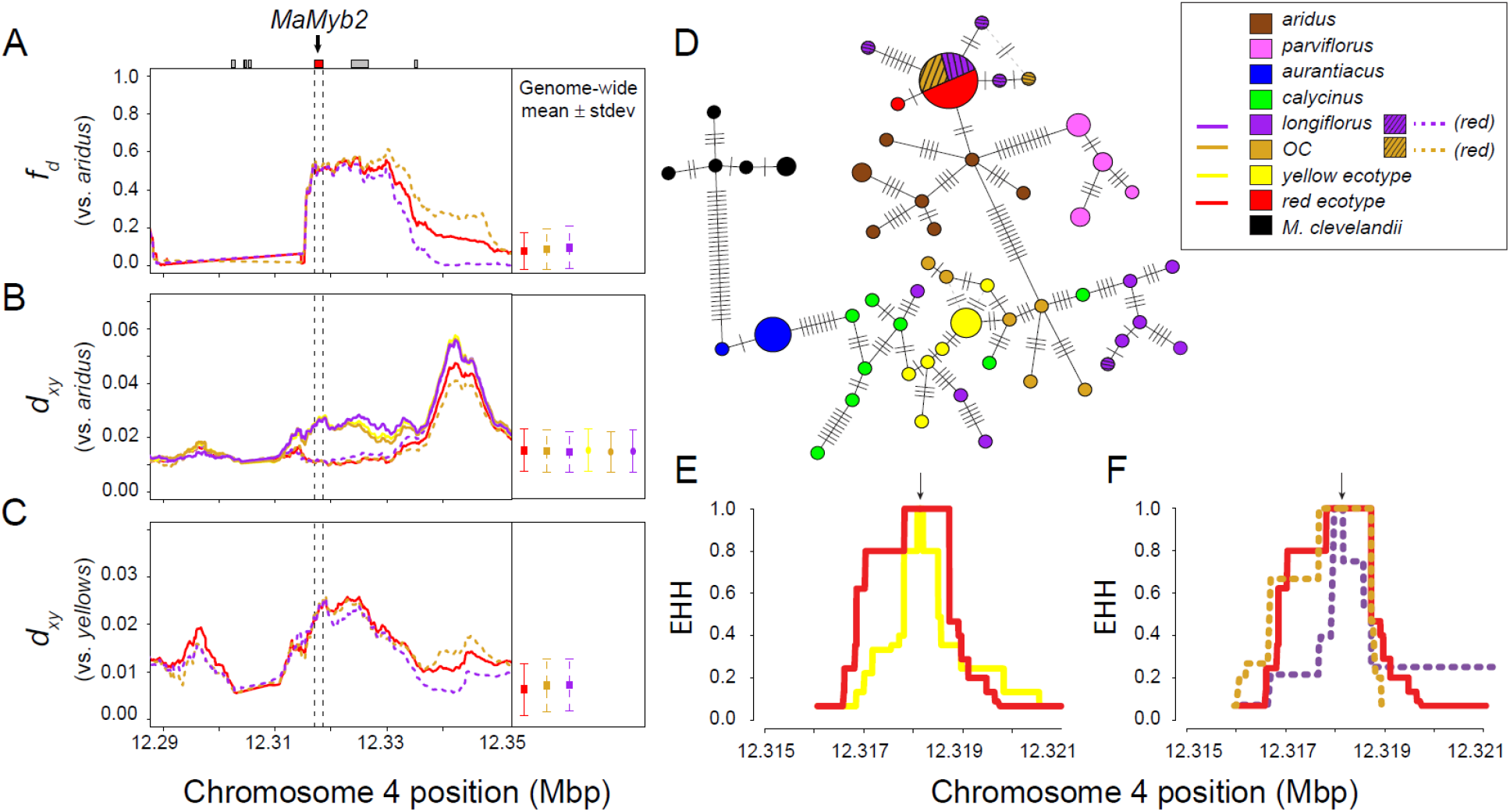
Ancient hybridization leads to the repeated origin of red flowers. (A-C) On the left, the admixture proportion (*f*_*d*_) and genetic divergence (*d*_*xy*_) are plotted across a 60 kb region surrounding the *MaMyb2* gene on chromosome 4. The dashed vertical line represents the start and stop positions of *MaMyb2*, and gray boxes at the top correspond to the position of additional genes in this region. On the right of each plot is the genome-wide mean and standard deviation of each statistic, calculated in 10kb windows and 100 bp steps across all 10 chromosomes. (A) *f*_*d*_ is calculated using the red-flowered samples from clade D as P2 (red ecotype, red-*OC*, and red-*longiflorus*), *aurantiacus* as P1, and *aridus* as P3. (B) *d*_*xy*_ between the red-flowered samples and *aridus* (dotted lines) is low across the *MaMyb2* region, but *d*_*xy*_ between *aridus* and each of their closely related yellow samples (yellow ecotype, yellow-*OC*, and yellow *longiflorus*) is elevated in this same region (solid lines). (C) *d*_*xy*_ calculated between the three sets of red-flowered samples and their most closely related yellow-flowered partners is elevated across this same region compared to genome-wide averages. (D) A haplotype network showing the number of sequence differences among unique *MaMyb2* haplotypes in all taxa. Each haplotype is represented by a circle, the size of which is proportional to its observed frequency. Black dashes show the number of mutational steps separating haplotypes. Alternate connections among haplotypes are shown as dashed, gray bars. (E) Site–specific extended haplotype homozygosity (EHH) for a site in the middle of *MaMyb2* (black arrow) decays more slowly for the red ecotype compared to the yellow ecotype. (F) EHH also decays more slowly for the red ecotype and red-*OC* samples than for the red-*longiflous* samples.

To demonstrate that the red-flowered taxa share a common ancestral sequence at *MaMyb2*, we compared haplotype variation across the entire *MaMyb2* gene in members of the radiation (Fig 5D). Consistent with findings from Stankowski and Streisfeld (2015), all 12 of the red-flowered individuals from clade D included in this study had shared sequences that grouped together with the distantly related *aridus*. Indeed, there was a single common haplotype that contained 9 of 10 sequenced chromosomes from the red ecotype, 5 of 6 red *OC* chromosomes, and 4 of 8 red *longiflorus* chromosomes. Moreover, this haplotype was separated by only two mutational steps to the closest *aridus* haplotype. By contrast, all yellow-flowered samples from clades C and D grouped together but were positioned distantly from the red-flowered haplotypes. The only exception was a single red-flowered *longiflorus* individual. While one of the two chromosomes grouped with the red-flowered samples, the other chromosome from this individual grouped with haplotypes from yellow *longiflorus*, implying that despite having red flowers, this individual was heterozygous for multiple SNPs within *MaMyb2*. Finally, the haplotypes identified in the red-flowered samples from clade D were more similar to haplotypes in the yellow-flowered *aridus* than those from its sister subspecies, the red-flowered *parviflorus*.

Despite the red ecotype and *parviflorus* sharing a common genetic basis for red flowers (Stankowski and Streisfeld 2015), the current data reveal genome-wide, heterogeneous signatures of introgression between *aridus* and members of clade D. Moreover, the *MaMyb2* sequences in the red-flowered taxa are more similar to *aridus* than they are to the red-flowered *parviflorus*, indicating that the introgressed block was donated from an ancestor of *aridus* after the split with *parviflorus*. These findings rule out two possibilities for the origins of red flowers in clade D: 1) that the red allele was introgressed from the ancestor of *aridus* and *parviflorus* into the ancestor of clade D, and 2) that the red allele at *MaMyb2* represents an ancient polymorphism that formed prior to the divergence between clades B and D and has sorted differentially into descendant lineages. It is also unlikely that *MaMyb2* was recently introgressed between *aridus* and one of the red-flowered taxa followed by ongoing gene flow among populations. The red-flowered populations are largely geographically isolated from each other and are surrounded by yellow-flowered populations, but no evidence of introgression was found in this genomic region in any of the yellow-flowered individuals we sequenced. Moreover, we find that sequence divergence at *MaMyb2* between the red-flowered samples is lower than divergence between the reds and *aridus*, and it is similar to levels of divergence between red-flowered plants across the genome (Fig. S3), suggesting that introgression in this region did not occur recently. Thus, the most likely explanation for the similarity between *MaMyb2* haplotypes from red-flowered samples and yellow-flowered *aridus* is that red-flowered variants present in ancestral populations of *aridus* hybridized with the ancestor of clade D but have since gone extinct. Although distinguishing between this hypothesis and others for the origins of red flowers will ultimately require finding the causal variant(s) responsible for red flowers in each of these taxa, the current results deepen our understanding of the origins of a haplotype associated with divergence and the early stages of speciation. Specifically, this haplotype originated in an ancestor of *aridus*, was shared with an ancestor of clade D, and has since been maintained in some populations and lost in others.

An additional question related to the maintenance of red flowers in these lineages is whether this shared haplotype shows consistent evidence of natural selection among taxa. Multiple lines of evidence support the presence of geographically widespread and recent positive selection in the red ecotype in San Diego. Specifically, despite ongoing gene flow between the ecotypes, there is a steep geographic cline in both allele frequency and ancestry at this locus (Streisfeld et al 2013; Stankowski et al 2023), and this locus shows a unique pattern of divergence relative to the rest of the genome (Stankowski et al 2017; 2019). However, it is unclear whether the other red-flowered samples also show evidence of positive selection following introgression.

We explored patterns of recent positive selection in the red-flowered taxa by calculating the site-specific extended haplotype homozygosity (EHH) statistic. Due to a recent selective sweep, a rapid increase in the frequency of a beneficial mutation will result in elevated linkage disequilibrium, leading to extended patterns of homozygosity in haplotypes (Smith and Haigh 1974, Sabeti et al. 2002, Voight et al. 2006). However, in the absence of selection, we expect haplotypes to break down over time due to new mutations and recombination. Indeed, we detected a broader region of EHH surrounding the *MaMyb2* gene in the red ecotype relative to the yellow ecotype (Fig. 5E). Individuals from populations of red-flowered *OC* have similar EHH to the red ecotype, but the haplotypes from red *longiflorus* are considerably shorter (Fig. 5F). These findings are consistent with a similar strength or timing of natural selection in the two *puniceus* red-flowered taxa, but the shorter haplotypes in red *longiflorus* suggest that consistent, directional selection on red flowers is weak, absent, or occurred deeper in the past in these populations relative to the red-flowered populations of *puniceus*. Although it is possible this red-flowered variant in *longiflorus* may be lost in the future, as may have been the case in the numerous populations across southern California that only produce yellow flowers, some form of fluctuating or frequency-dependent selection also may maintain the different flower color variants in these polymorphic populations (Schemske and Bierzychudek 2001). Future work directly connecting the effects of flower color with fitness differences in the wild will be necessary to evaluate these alternate hypotheses. Regardless, these findings make it clear that the ancient introduction of this haplotype into the ancestor of clade D has had important consequences for the evolution of floral diversity in this group.

## Conclusions

The results from this study demonstrate how hybridization can occur early in the history of a radiation, but that widespread selection against gene flow can reduce that signal over time. However, hybridization also can introduce beneficial variation that promotes divergence by facilitating adaptation to new ecological niches. The introgression of the *MaMyb2* gene early in the divergence of clade D has fueled the repeated evolution of red flowers in this radiation, which has led to a barrier to gene flow between the red and yellow ecotypes. In addition, there also appear to be numerous barriers to gene flow in place between *aridus* and members of clade D, which likely limit ongoing hybridization between these taxa in nature. More broadly, the re-use of old genetic variation has become a leading explanation for rapid diversification in evolutionary radiations (Marques et al 2019). In some cases, introgression was not the source of this variation, as in threespine sticklebacks that repeatedly colonized freshwater lakes due to the presence of extensive standing variation in the ancestral, marine population (Nelson and Cresko 2018). Alternatively, radiations of cichlid fish and Hawaiian silverswords appear to have benefited directly from ancient hybridization, which led to the remarkable morphological diversity we see today in those groups (Barrier et al. 1999, Meier et al. 2017). In *Mimulus*, we have shown that recent natural selection has preserved introgressed haplotypes in the red-flowered taxa, which have been lost in yellow-flowered plants. Thus, hybridization can be a creative force in evolution, but low fitness in hybrids also can lead to the evolution of reproductive barriers, both of which reveal gene flow’s important role in promoting and maintaining diversity.

## Materials and Methods

### Genome resequencing and variant calling

Leaf tissue was collected from the wild (Table S1), and DNA from the 10 new samples was extracted, prepared, and sequenced as described in Stankowski et al. (2019). Variant calling, filtering, and haplotype phasing was performed as described in Stankowski et al. (2019) after mapping reads from all 47 samples to the *M. aurantiacus* reference genome. We then further filtered the VCF file for biallelic SNPs and removed sites with missing genotype data. The final data set contained 12,730,568 SNPs across all 47 samples.

### PCAs and Network analysis

We generated a neighbor-joining splits network using *SplitsTree4 v*.*4*.*17*.1 (Huson, Kloepper, & Bryant 2008) from all 47 samples. We filtered SNPs for linkage disequilibrium in *Plink* version 1.90 (Chang et al. 2015) to remove variant sites with an *r*^*2*^ greater than 0.2 in 50 kb windows, sliding by 10 kb, resulting in a data set containing 1,640,850 SNPs. To assess clustering patterns among more closely related samples, we ran a principal components analysis (PCA) in *Plink* using only the 27 samples from clade D.

### Tests for genome-wide admixture

We calculated Patterson’s *D* for all possible groups of three in-group taxa using the Dtrios command in the *Dsuite* software package (Malinsky, Matschiner, & Svardal 2021) based on the relationships inferred from genome-wide data in Stankowski et al. (2019), with *M. clevelandii* used as the out-group for all tests. The four samples from subspecies *grandiflorus* were removed prior to this analysis, as the calculation of *D* using these samples does not provide information about the history of introgression between clade D and *aridus*. We corrected for multiple tests using a Bonferroni-corrected alpha of 0.0009.

*f*_*d*_ was calculated in 50kb non-overlapping genomic windows using the ABBABABAwindows.py Python script (https://github.com/simonhmartin/genomics_general) and smoothed across the genome using the *loess* function from the *scales* package (Wickham and Seidel 2022) in R with a span of 0.02. To determine the average level of admixed genetic variation between *aridus* and the yellow ecotype for P1 taxa at different levels of taxonomic divergence, we estimated the mean *f*_*d*_ from each test across 50 kb windows. To estimate the average level of sequence divergence between the yellow ecotype and the taxa from clades C and D, we calculated average *d*_*a*_ in 50 kb windows for each pair of taxa (Nei & Li 1979). *d*_*a*_ accounts for genetic divergence that predated species divergence by subtracting the average intraspecific pairwise differences (π in both species) from the observed interspecific value (*d*_*xy*_). π and *d*_*xy*_ were calculated using PIXY version 1.2.5 (Korunes & Samuk 2021) with both variant and invariant sites included. To test for significant differences among the mean *f*_*d*_ values, we fit a linear model with *f*_*d*_ as the dependent variable and the P1 taxon as the independent variable using the *lm* function in R and then used the *emmeans* package (Lenth, 2019) to perform pairwise comparisons of the estimated marginal mean *f*_*d*_ values for the different P1s from the linear model.

We estimated the Spearman’s rank correlation coefficient between *f*_*d*_, *F*_*ST*_, and π using the *cor*.*test* function in R. *F*_*ST*_ was calculated in each 50 kb window between the yellow ecotype and one of the five remaining taxa from clades C and D using the *popgenwindows*.*py* Python script (https://github.com/simonhmartin/genomics_general), with only variant sites included. Finally, we sorted the *f*_*d*_ values calculated in 50 kb windows into quantile bins based on recombination rates estimated in 500kb windows in Stankowski et al. (2019). We then calculated the mean and 95% confidence intervals of the *f*_*d*_ values within each recombination rate quantile bin. To test for differences in *f*_*d*_ among recombination bins, we fit a linear model with *f*_*d*_ as the dependent variable and recombination bin as the independent variable and then perform pairwise comparisons of the estimated marginal mean *f*_*d*_ values for the different recombination bins from the linear model.

### Admixture and genetic divergence across the *MaMyb2* region

To determine the history of the red allele at *MaMyb2* in the different lineages of clade D, we calculated *f*_*d*_ and *d*_*xy*_ as described above, but in overlapping 10 kb windows with 100 bp steps. *f*_*d*_ was calculated three times, each time with *aurantiacus* as P1 and *aridus* as P3, but varying the different red samples (red ecotype, red-OC, and red *longiflorus*) as P2. *d*_*xy*_ was calculated between each of the red taxa and *aridus*, each of their closest yellow-flowered partners and *aridus*, between each of the red-flowered taxa, and between each of the red taxa and their most closely related yellow partners.

### Haplotype Network

We used VCFx version: 2.0.6b to generate a FASTA file of the entire *MaMyb2* gene (positions 12,317,113 to 12,318,500 on chromosome 4) from the VCF file from all 47 samples. This FASTA file was then used to construct a haplotype network from all recovered haplotypes based on an infinite site model and uncorrected distances using *Pegas* version 1.1 (Paradis 2010) in R version 3.6.3.

### Extended Haplotype Homozygosity

We calculated the site-specific extended haplotype homozygosity statistic for a SNP in the middle of the *MaMyb2* gene (position 12,317,808) using *rehh* version 3.2.2 (Gautier & Vitalis 2012). Separate VCFs were created that contained samples from the red ecotype, the yellow ecotype, red-*OC*, or red*-longiflorus*. Haplotype and marker information was extracted from each VCF file using the *data2haplohh* function of the *rehh* package (Gautier & Vitalis 2012). EHH was calculated from the marker and haplotype information for each of the taxa using the *calc_ehh* function in *rehh* (Gautier & Vitalis 2012).

## Data Availability

Raw sequencing reads were downloaded from the Short-Read Archive (SRA) from bioproject ID: PRJNA549183. New sequencing reads generated here are included as bioproject ID PRJNAXXXXX. VCF files and population genomic data have been deposited on DRYAD (link to come). The reference assembly and annotation are available at mimubase.org. Computer scripts used for population genomic analysis are available on Github at: https://github.com/awshort/aridus_introgression.

## Acknowledgments

We would like to thank Sean Stankowski and Madeline Chase for their help in generating the Illumina libraries and motivating this project. We would also like to thank Peter Ralph, Andrew Kern, Yaniv Brandvain, and Bill Cresko for providing valuable feedback and discussion. Doug Turnbull and Maggie Weitzman conducted the Illumina sequencing at the UO Genomics & Cell Characterization Core Facility (GC3F).

