## supplement for "Ancient hybridization leads to the repeated evolution of red flowers across a monkeyflower radiation"

Table S1. Sample information for the 47 sequenced individuals used in this study. The table includes taxon identity, sampling location, percent read alignment, and average sequencing depth. Samples in red text were sequenced as a part of this study, while those in black were described in Stankowski et al (2019).

| Sample | Taxon | Latitude | Longitude | % Reads aligned | Seq. Depth |
|---|---|---|---|---|---|
| **159_83** | ssp. *aridus* | 32.6630 | -116.2230 | 91.7 | 21.12 |
| **159_84** | ssp. *aridus* | 32.6630 | -116.2230 | 89.3 | 21.98 |
| **195_1** | ssp. *aridus* | 32.6300 | -116.1429 | 92.6 | 20.20 |
| **T84** | ssp. *aridus* | 32.6526 | -116.2449 | 87.2 | 21.75 |
| **T102** | ssp. *aurantiacus* | 39.0424 | -122.7727 | 94.9 | 23.74 |
| **T104** | ssp. *aurantiacus* | 39.2045 | -123.7646 | 94.6 | 25.09 |
| **T50** | ssp. *aurantiacus* | 35.9865 | -121.4928 | 88.3 | 24.36 |
| **T92** | ssp. *aurantiacus* | 37.8459 | -120.6110 | 94.0 | 15.16 |
| **T144** | ssp. *calycinus* | 34.1929 | -117.2784 | 93.2 | 26.00 |
| **T150** | ssp. *calycinus* | 33.8564 | -116.8481 | 94.7 | 24.02 |
| **T90** | ssp. *calycinus* | 35.5918 | -118.5052 | 91.3 | 19.97 |
| **T91** | ssp. *calycinus* | 35.3172 | -118.5871 | 95.5 | 27.91 |
| **T101** | ssp. *grandiflorus* | 39.5536 | -121.4301 | 92.0 | 16.05 |
| **T61** | ssp. *grandiflorus* | 39.5590 | -120.8243 | 91.6 | 17.31 |
| **T96** | ssp. *grandiflorus* | 39.0122 | -120.7552 | 92.0 | 28.21 |
| **T99** | ssp. *grandiflorus* | 39.4376 | -121.0599 | 91.4 | 23.84 |
| **DPR_Y3** | ssp. *longiflorus*, yellow | 33.7459 | -117.4485 | 96.0 | 26.88 |
| **SS_Y16** | ssp. *longiflorus*, yellow | 34.2722 | -118.6100 | 94.2 | 30.86 |
| **T33_1** | ssp. *longiflorus*, yellow | 34.3438 | -118.5099 | 94.6 | 18.87 |
| **T8_8** | ssp. *longiflorus*, yellow | 34.1347 | -118.6452 | 82.6 | 25.11 |
| **DPR_R12** | ssp. *longiflorus*, red | 33.7459 | -117.4485 | 89.8 | 18.52 |
| **MBR_R3** | ssp. *longiflorus*, red | 34.2056 | -117.6765 | 94.4 | 23.20 |
| **SS_R19** | ssp. *longiflorus*, red | 34.2722 | -118.6100 | 91.3 | 23.89 |
| **WTF_R1** | ssp. *longiflorus*, red | 34.2174 | -117.7630 | 94.2 | 23.45 |
| **KK168** | ssp. *parviflorus* | 34.0180 | -119.6730 | 91.8 | 23.66 |
| **KK161** | ssp. *parviflorus* | 34.0180 | -119.6730 | 92.0 | 19.11 |
| **KK180** | ssp. *parviflorus* | 34.0180 | -119.6730 | 92.4 | 18.18 |
| **KK182** | ssp. *parviflorus* | 34.0193 | -119.6802 | 91.3 | 19.46 |
| **MTH17** | ssp. *puniceus*, *OC*, red | 33.6414 | -117.8252 | 90.6 | 21.23 |
| **MTH15** | ssp. *puniceus*, *OC*, red | 33.6414 | -117.8252 | 91.1 | 18.85 |
| **RPR16** | ssp. *puniceus*, *OC*, red | 33.6058 | -117.8020 | 95.1 | 21.68 |
| **LFP13** | ssp. *puniceus*, *OC*, yellow | 33.6532 | -117.6579 | 90.3 | 22.23 |
| **VCR14** | ssp. *puniceus*, *OC*, yellow | 33.4871 | -117.6489 | 92.2 | 19.97 |
| **VCR17** | ssp. *puniceus*, *OC*, yellow | 33.4871 | -117.6489 | 90.7 | 21.38 |
| **ELF** | ssp. *puniceus*, red | 33.0860 | -117.1453 | 93.0 | 18.20 |

| | | | | | |
|---|---|---|---|---|---|
| **JMC** | ssp. *puniceus*, red | 32.7373 | -116.9541 | 93.8 | 19.06 |
| **LH** | ssp. *puniceus*, red | 33.0609 | -117.1188 | 87.1 | 19.77 |
| **MT** | ssp. *puniceus*, red | 32.8210 | -117.0618 | 93.7 | 20.85 |
| **UCSD** | ssp. *puniceus*, red | 32.8894 | -117.2362 | 87.0 | 18.23 |
| **BCRD** | ssp. *puniceus*, yellow | 32.9496 | -116.6380 | 94.6 | 20.85 |
| **INJ** | ssp. *puniceus*, yellow | 33.0979 | -116.6643 | 93.1 | 18.83 |
| **LO** | ssp. *puniceus*, yellow | 32.6767 | -116.3312 | 93.4 | 18.04 |
| **PCT** | ssp. *puniceus*, yellow | 32.7326 | -116.4698 | 92.3 | 19.68 |
| **POTR** | ssp. *puniceus*, yellow | 32.6038 | -116.6339 | 90.5 | 19.27 |
| **CLV_GH** | *M. clevelandii* | 33.1589 | -116.8122 | 92.3 | 21.31 |
| **CLV_11** | *M. clevelandii* | 33.3391 | -116.9325 | 84.4 | 15.52 |
| **CLV_4** | *M. clevelandii* | 33.3391 | -116.9325 | 89.3 | 17.31 |

Table S2. Patterson's D statistics, estimated using the *Dtrios* function in the *Dsuite* package, for all trios of taxa across the *M. aurantiacus* species complex. In all cases, *M. clevelandii* is used as the outgroup, and the three ingroup taxa are indicated (P1, P2, P3). The identify of the P1 and P2 taxa is adjusted, so that the D-statistics are always positive. Counts of each site category (BBAA, ABBA, and BABA) are indicated.

| P1 | P2 | P3 | Dstatistic | Z-score | p-value | f4-ratio | BBAA | ABBA | BABA |
|---|---|---|---|---|---|---|---|---|---|
| *aurantiacus* | *calycinus* | *aridus* | 0.0247 | 3.5809 | 3.4236E-04 | 0.0085 | 503634 | 144599 | 137624 |
| *longiflorus* | *calycinus* | *aridus* | 0.0231 | 5.6748 | 1.3888E-08 | 0.0066 | 585489 | 120190 | 114762 |
| *aurantiacus* | *longiflorus* | *aridus* | 0.0055 | 0.7555 | 4.4995E-01 | 0.0019 | 500695 | 140659 | 139113 |
| *aurantiacus* | *OC* | *aridus* | 0.0393 | 3.4859 | 4.9054E-04 | 0.0140 | 484366 | 152419 | 140898 |
| *calycinus* | *OC* | *aridus* | 0.0169 | 2.0655 | 3.8877E-02 | 0.0056 | 531413 | 136476 | 131930 |
| *longiflorus* | *OC* | *aridus* | 0.0389 | 4.2424 | 2.2114E-05 | 0.0122 | 545247 | 133213 | 123239 |
| *aurantiacus* | *Red ecotype* | *aridus* | 0.0720 | 5.5712 | 2.5299E-08 | 0.0264 | 476747 | 161375 | 139700 |
| *calycinus* | *Red ecotype* | *aridus* | 0.0524 | 4.8891 | 1.0132E-06 | 0.0180 | 515736 | 147655 | 132955 |
| *longiflorus* | *Red ecotype* | *aridus* | 0.0743 | 6.3053 | 2.8760E-10 | 0.0245 | 525314 | 145558 | 125430 |
| *OC* | *Red ecotype* | *aridus* | 0.0440 | 6.5784 | 4.7556E-11 | 0.0125 | 590243 | 120491 | 110336 |
| *aurantiacus* | *Yellow ecotype* | *aridus* | 0.0976 | 7.9287 | 2.2138E-15 | 0.0366 | 472730 | 169291 | 139178 |
| *calycinus* | *Yellow ecotype* | *aridus* | 0.0817 | 7.9242 | 2.2968E-15 | 0.0284 | 517873 | 153126 | 129988 |
| *longiflorus* | *Yellow ecotype* | *aridus* | 0.1033 | 9.5022 | 0.0000E+00 | 0.0348 | 524357 | 152595 | 124029 |
| *OC* | *Yellow ecotype* | *aridus* | 0.0741 | 11.6458 | 0.0000E+00 | 0.0229 | 563239 | 134736 | 116144 |
| *Red ecotype* | *Yellow ecotype* | *aridus* | 0.0354 | 5.3659 | 8.0570E-08 | 0.0105 | 586759 | 123454 | 115016 |
| *longiflorus* | *calycinus* | *aurantiacus* | 0.0142 | 3.0375 | 2.3856E-03 | 0.0152 | 264814 | 158551 | 154123 |
| *OC* | *calycinus* | *aurantiacus* | 0.0646 | 12.9041 | 0.0000E+00 | 0.0728 | 226218 | 185771 | 163228 |
| *Red ecotype* | *calycinus* | *aurantiacus* | 0.0810 | 13.8921 | 0.0000E+00 | 0.0916 | 217043 | 193298 | 164335 |
| *Yellow ecotype* | *calycinus* | *aurantiacus* | 0.0914 | 16.1788 | 0.0000E+00 | 0.1016 | 222638 | 193789 | 161330 |
| *OC* | *longiflorus* | *aurantiacus* | 0.0542 | 7.6557 | 1.9227E-14 | 0.0585 | 238017 | 176044 | 157929 |
| *Red ecotype* | *longiflorus* | *aurantiacus* | 0.0710 | 9.0906 | 1.0842E-19 | 0.0776 | 224813 | 184963 | 160429 |
| *Yellow ecotype* | *longiflorus* | *aurantiacus* | 0.0810 | 12.6167 | 0.0000E+00 | 0.0878 | 227424 | 187130 | 159100 |
| *aridus* | *parviflorus* | *aurantiacus* | 0.1961 | 14.0213 | 0.0000E+00 | 0.1348 | 285314 | 268360 | 180373 |
| *Red ecotype* | *OC* | *aurantiacus* | 0.0230 | 4.3617 | 1.2905E-05 | 0.0203 | 290560 | 142601 | 136181 |
| *Yellow ecotype* | *OC* | *aurantiacus* | 0.0330 | 4.9649 | 6.8722E-07 | 0.0310 | 270354 | 155205 | 145289 |
| *Yellow ecotype* | *Red ecotype* | *aurantiacus* | 0.0126 | 2.2007 | 2.7757E-02 | 0.0109 | 297206 | 140834 | 137339 |
| *aridus* | *parviflorus* | *calycinus* | 0.1748 | 10.9417 | 0.0000E+00 | 0.1579 | 285807 | 267415 | 187841 |
| *Red ecotype* | *OC* | *calycinus* | 0.0555 | 9.6966 | 0.0000E+00 | 0.1384 | 243727 | 158758 | 142055 |
| *Yellow ecotype* | *OC* | *calycinus* | 0.0356 | 6.1916 | 5.9553E-10 | 0.1002 | 220904 | 168746 | 157148 |
| *Red ecotype* | *Yellow ecotype* | *calycinus* | 0.0171 | 3.8015 | 1.4382E-04 | 0.0421 | 250570 | 152011 | 146907 |
| *aridus* | *parviflorus* | *longiflorus* | 0.1850 | 12.9018 | 0.0000E+00 | 0.1572 | 287491 | 267691 | 184097 |
| *Red ecotype* | *OC* | *longiflorus* | 0.0723 | 14.5433 | 0.0000E+00 | 0.1678 | 231928 | 164056 | 141932 |
| *Yellow ecotype* | *OC* | *longiflorus* | 0.0656 | 13.0470 | 0.0000E+00 | 0.1650 | 211269 | 176209 | 154529 |
| *Red ecotype* | *Yellow ecotype* | *longiflorus* | 0.0015 | 0.3270 | 7.4369E-01 | 0.0034 | 242556 | 151013 | 150569 |
| *calycinus* | *aurantiacus* | *parviflorus* | 0.0048 | 0.6611 | 5.0857E-01 | 0.0024 | 428154 | 150130 | 148692 |
| *longiflorus* | *aurantiacus* | *parviflorus* | 0.0095 | 1.0433 | 2.9681E-01 | 0.0047 | 425178 | 151583 | 148737 |
| *OC* | *aurantiacus* | *parviflorus* | 0.0213 | 2.2775 | 2.2757E-02 | 0.0107 | 412604 | 157123 | 150556 |
| *Red ecotype* | *aurantiacus* | *parviflorus* | 0.0214 | 2.5212 | 1.1695E-02 | 0.0108 | 406985 | 157924 | 151306 |
| *Yellow ecotype* | *aurantiacus* | *parviflorus* | 0.0199 | 2.3536 | 1.8592E-02 | 0.0102 | 405744 | 160179 | 153933 |
| *longiflorus* | *calycinus* | *parviflorus* | 0.0057 | 0.9576 | 3.3825E-01 | 0.0023 | 509818 | 124093 | 122685 |
| *OC* | *calycinus* | *parviflorus* | 0.0186 | 2.9785 | 2.8963E-03 | 0.0084 | 460163 | 140254 | 135125 |
| *Red ecotype* | *calycinus* | *parviflorus* | 0.0183 | 3.2225 | 1.2706E-03 | 0.0084 | 447078 | 143871 | 138691 |
| *Yellow ecotype* | *calycinus* | *parviflorus* | 0.0170 | 3.2455 | 1.1726E-03 | 0.0078 | 451949 | 143638 | 138830 |
| *OC* | *longiflorus* | *parviflorus* | 0.0139 | 1.8662 | 6.2017E-02 | 0.0061 | 473694 | 135280 | 131558 |
| *Red ecotype* | *longiflorus* | *parviflorus* | 0.0137 | 2.5911 | 9.5673E-03 | 0.0061 | 456355 | 140065 | 136292 |
| *Yellow ecotype* | *longiflorus* | *parviflorus* | 0.0122 | 2.2038 | 2.7536E-02 | 0.0055 | 457988 | 141254 | 137854 |
| *Red ecotype* | *OC* | *parviflorus* | 0.0002 | 0.0429 | 9.6576E-01 | 0.0001 | 524946 | 114939 | 114888 |
| *aurantiacus* | *Yellow ecotype* | *parviflorus* | -0.0199 | 2.3536 | 1.8592E-02 | 0.0102 | 405744 | 160179 | 153933 |
| *calycinus* | *Yellow ecotype* | *parviflorus* | -0.0170 | 3.2455 | 1.1726E-03 | 0.0078 | 451949 | 143638 | 138830 |
| *longiflorus* | *Yellow ecotype* | *parviflorus* | -0.0122 | 2.2038 | 2.7536E-02 | 0.0055 | 457988 | 141254 | 137854 |
| *OC* | *Yellow ecotype* | *parviflorus* | 0.0013 | 0.2585 | 7.9602E-01 | 0.0005 | 500637 | 123762 | 123441 |
| *OC* | *Yellow ecotype* | *parviflorus* | 0.0013 | 0.2585 | 7.9602E-01 | 0.0005 | 500637 | 123762 | 123441 |
| *Red ecotype* | *Yellow ecotype* | *parviflorus* | 0.0016 | 0.6136 | 5.3950E-01 | 0.0006 | 527218 | 115540 | 115168 |
| *Red ecotype* | *Yellow ecotype* | *parviflorus* | 0.0016 | 0.6136 | 5.3950E-01 | 0.0006 | 527218 | 115540 | 115168 |
| *calycinus* | *longiflorus* | *OC* | 0.0521 | 6.8952 | 5.3776E-12 | 0.0921 | 221249 | 172531 | 155434 |
| *aridus* | *parviflorus* | *OC* | 0.1534 | 8.9739 | 3.2526E-19 | 0.1204 | 286312 | 262790 | 192892 |
| *Yellow ecotype* | *Red ecotype* | *OC* | 0.1006 | 13.8492 | 0.0000E+00 | 0.2457 | 181411 | 179420 | 146609 |
| *calycinus* | *longiflorus* | *Red ecotype* | 0.0368 | 4.7774 | 1.7761E-06 | 0.0725 | 235255 | 164413 | 152737 |
| *aridus* | *parviflorus* | *Red ecotype* | 0.1300 | 7.2320 | 4.7589E-13 | 0.1129 | 283071 | 259498 | 199805 |
| *calycinus* | *longiflorus* | *Yellow ecotype* | 0.0220 | 3.1743 | 1.5023E-03 | 0.0576 | 233650 | 163251 | 156236 |
| *aridus* | *parviflorus* | *Yellow ecotype* | 0.1114 | 6.1979 | 5.7217E-10 | 0.1062 | 280770 | 257570 | 205942 |

Table S3. The *t*-ratio from a linear mixed-effects model showing the difference in admixture proportion ($f_d$) using different taxa as P1, with the yellow ecotype as P2, *aridus* as P3 and *M. clevelandii* as the outgroup. Statistical significance is denoted as: *** for p < 0.001, ** for p < 0.01 and * for p < 0.05.

| **Factor Level** | **Estimate** | ***df*** | ***t*-ratio** |
|---|---|---|---|
| *aurantiacus* vs *calycinus* | 0.016113 | 11102 | 6.945*** |
| *aurantiacus* vs *longiflorus* | 0.015227 | 11102 | 6.65*** |
| *aurantiacus* vs *OC* | 0.025113 | 11102 | 10.757*** |
| *aurantiacus* vs red ecotype | 0.036214 | 11102 | 15.363*** |
| *calycinus* vs *longiflorus* | -0.000885 | 11102 | -0.387 |
| *calycinus* vs *OC* | 0.009 | 11102 | 3.862** |
| *calycinus* vs red ecotype | 0.020101 | 11102 | 8.542*** |
| *longiflorus* vs *OC* | 0.009885 | 11102 | 4.298*** |
| *longiflorus* vs *OC* | 0.020987 | 11102 | 9.033*** |
| *OC* vs red ecotype | 0.011102 | 11102 | 4.69*** |

Table S4. The *t*-ratio from a linear mixed-effects model showing the difference in admixture proportion ($f_d$) due to recombination rate. Quantile bins of recombination rate (in cM/Mb) are presented. Statistical significance is denoted as: *** for p < 0.001, ** for p < 0.01, and * for p < 0.05.

| **Factor Level** | **Estimate** | ***df*** | ***t*-ratio** |
|---|---|---|---|
| [0,0.824] vs (0.824,1.51] | -0.001286 | 10577 | -0.526 |
| [0,0.824] vs (1.51,2.33] | -0.001071 | 10577 | -0.446 |
| [0,0.824] vs (2.33,3.66] | -0.020672 | 10577 | -8.687*** |
| [0,0.824] vs (3.66,10.9] | -0.035771 | 10577 | -15.431*** |
| (0.824,1.51] vs (1.51,2.33] | 0.000215 | 10577 | 0.089 |
| (0.824,1.51] vs (2.33,3.66] | -0.019386 | 10577 | -8.089*** |
| (0.824,1.51] vs (3.66,10.9] | -0.034486 | 10577 | -14.766*** |
| (1.51,2.33] vs (2.33,3.66] | -0.019601 | 10577 | -8.33*** |
| (1.51,2.33] vs (3.66,10.9] | -0.034700 | 10577 | -15.149*** |
| (2.33,3.66] vs (3.66,10.9] | -0.015099 | 10577 | -6.654*** |

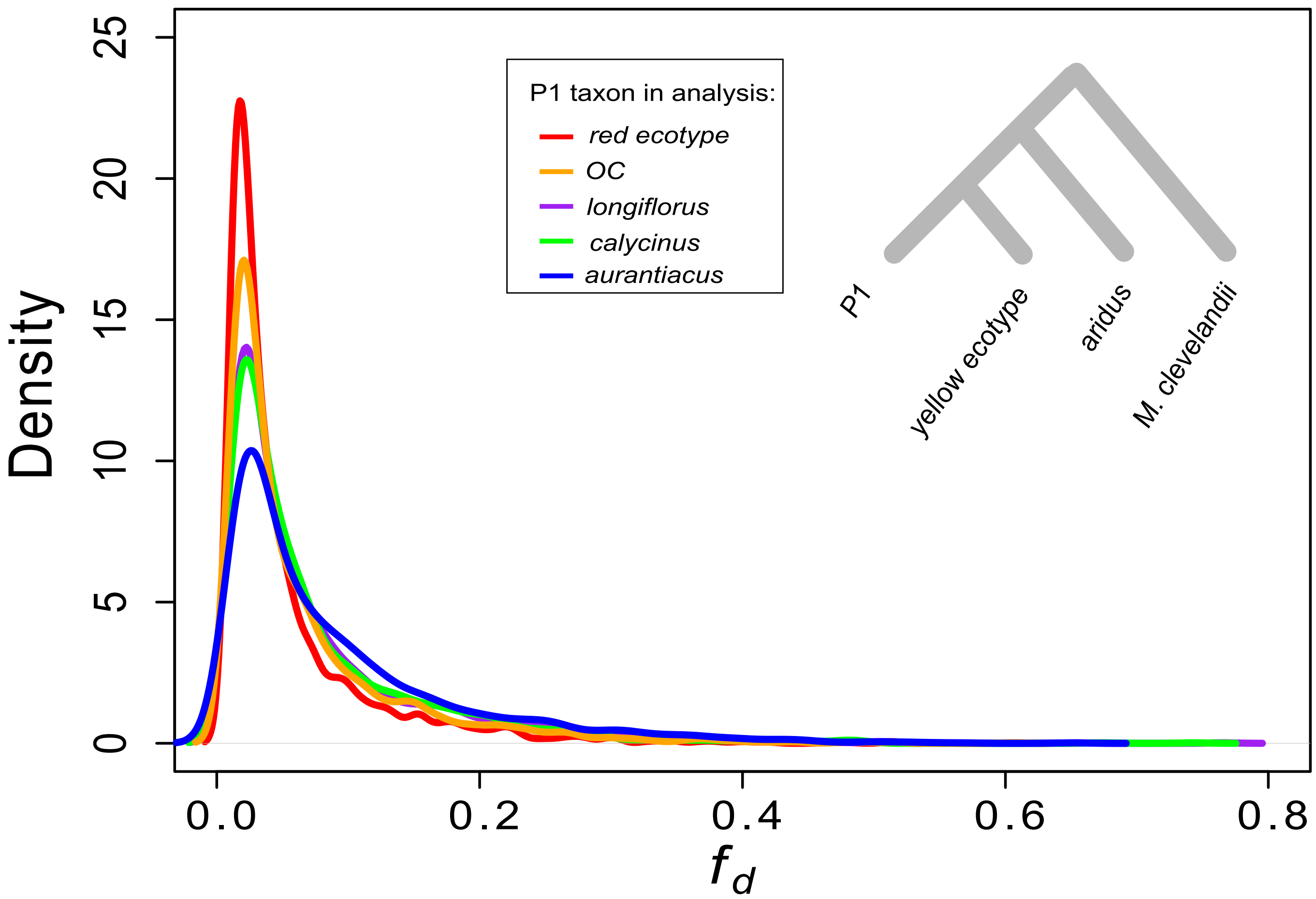


Fig. S1. Density plots of $f_d$, calculated in 50 kb windows across the genome. The P1 taxon varied in each test, with P2 set as the yellow ecotype, P3 set as *aridus*, and the outgroup is *M. clevelandii*.

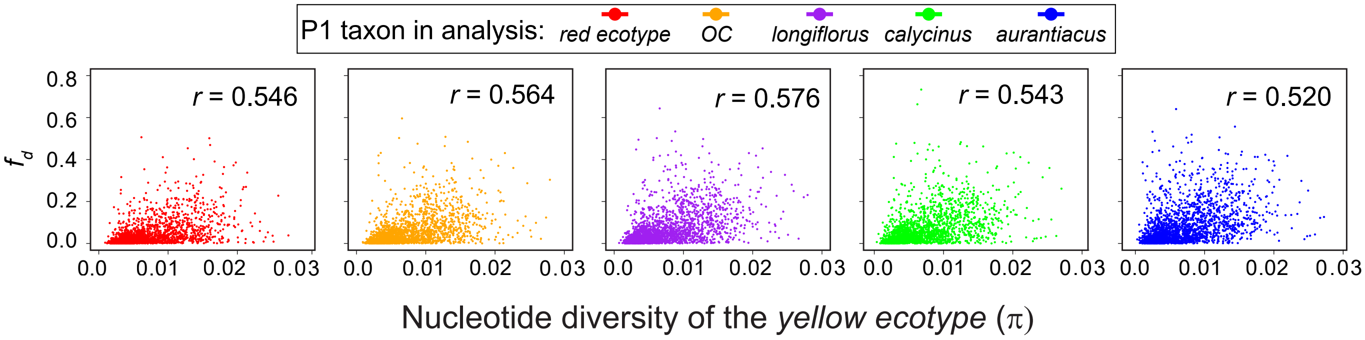


Fig. S2. Scatterplots and correlations between $f_d$ and π in 50 kb windows. $f_d$ is calculated with different P1 taxa, the yellow ecotype as P2, *aridus* as P3, and *M. clevelandii* as the outrgroup.

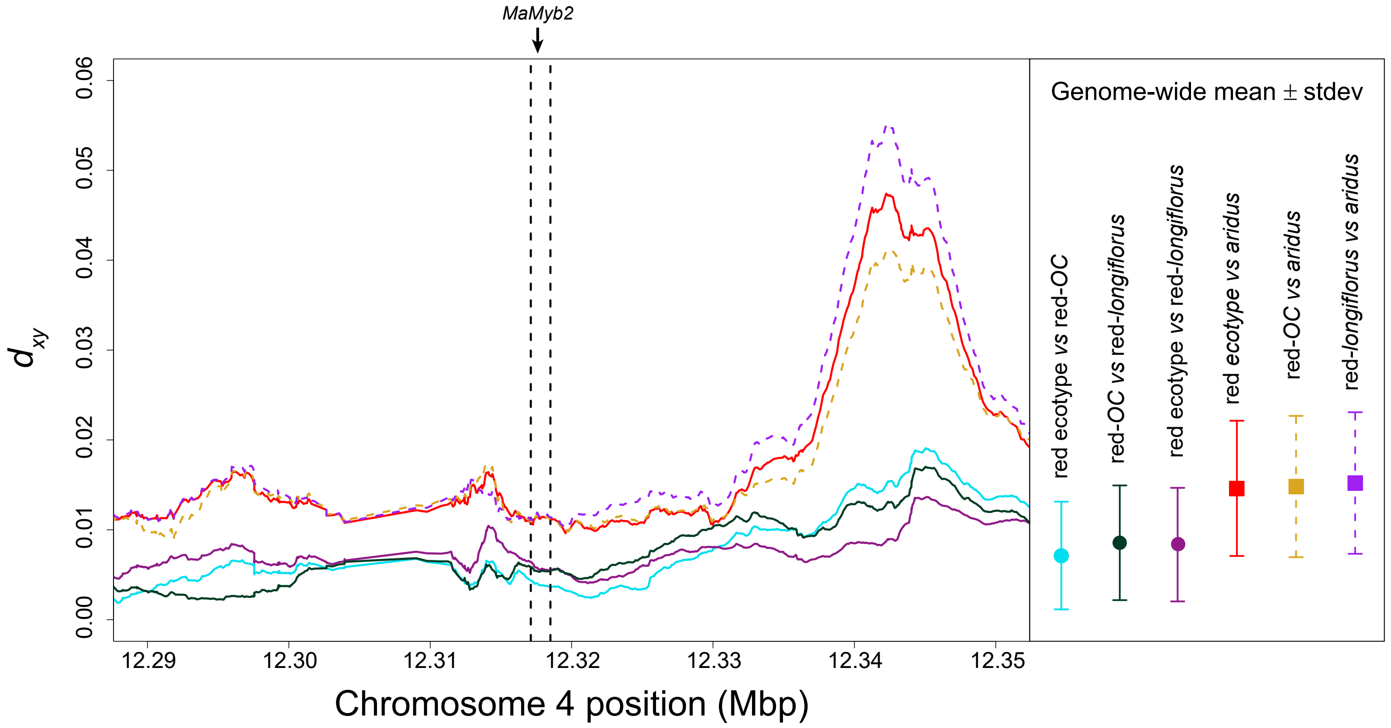


Fig. S3. Scans of $d_{xy}$ along chromosome 4, in the vicinity of *MaMyb2*. Divergence is calculated in overlapping 10 kb windows (100 bp steps) either between each of the three red-flowered taxa and *aridus*, or between each of the red-flowered taxa. The genome-wide mean values of $d_{xy}$ (error bars = standard deviations) are presented to the right, with the color scheme used the same as the scans.
